# Zika Virus Capsid Anchor Forms Cytotoxic Amyloid-like Fibrils

**DOI:** 10.1101/2020.11.13.381988

**Authors:** Kumar Udit Saumya, Kundlik Gadhave, Amit Kumar, Rajanish Giri

## Abstract

Capsid-anchor (CA) of Zika virus (ZIKV) is a small, single-pass transmembrane sequence that separates the capsid (C) protein from downstream pre-membrane (PrM) protein. During ZIKV polyprotein processing, CA is cleaved-off from C and PrM and left as a membrane-embedded peptide. CA plays an essential role in the assembly and maturation of the virus. However, its independent folding behavior is still unknown. Since misfolding and aggregation propensity of transmembrane proteins are now increasingly recognized and has been linked to several proteopathic disorders. Therefore, in this study, we investigated the amyloid-forming propensity of CA at physiological conditions. We observed aggregation behavior of CA peptide using dye-binding assays and ThT kinetics. The morphological analysis of CA aggregates explored by high-resolution microscopy (TEM and AFM) revealed characteristic amyloid-like fibrils. Further, the effect on mammalian cells exhibited the cytotoxic nature of the CA amyloid-fibrils. Our findings collectively shed light on the amyloidogenic phenomenon of flaviviral protein, which may contribute to their infection.

**Graphical Abstract:** Schematic representation of Zika virus Capsid anchor forming amyloid aggregates with cytotoxic and hemolytic properties.

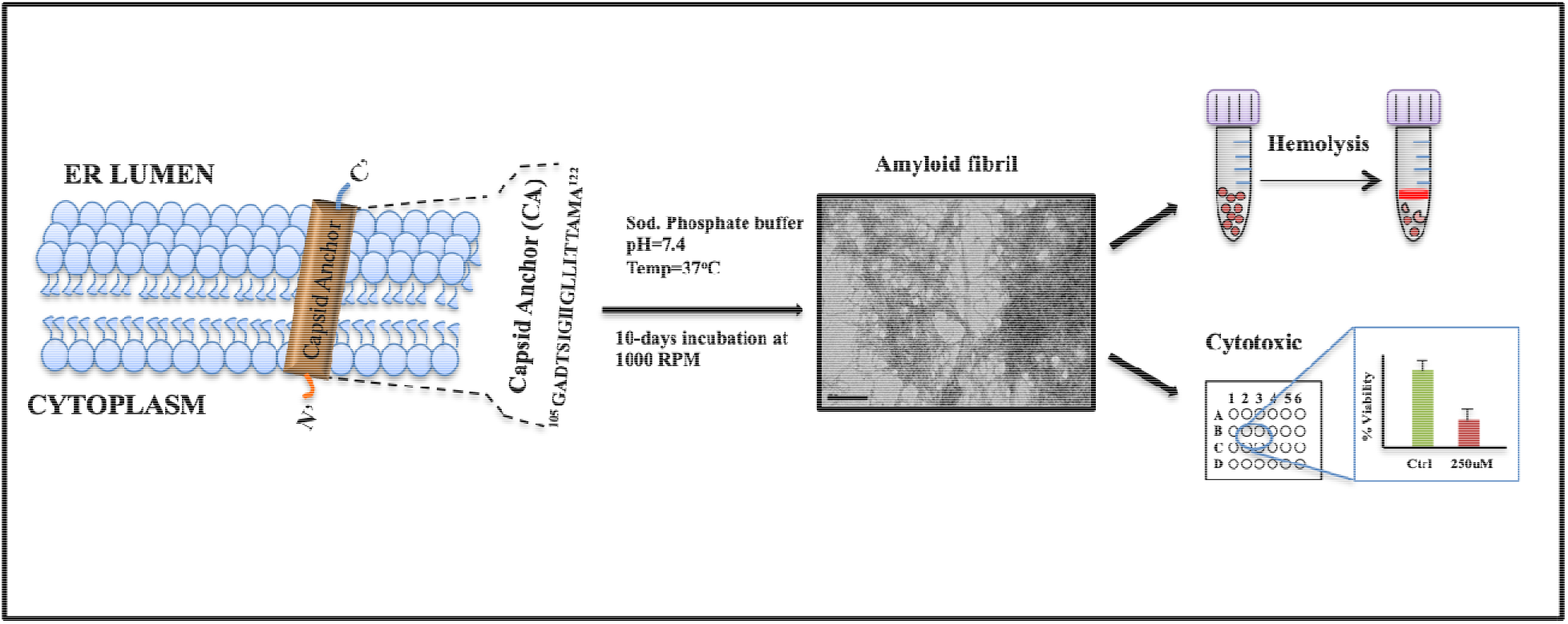

## Introduction

Zika virus (ZIKV) is an arthropod-borne flavivirus that was first isolated in the 1940’s from the Ugandan forest. In addition to transmission by mosquitoes, studies suggest this particular virus is also spread through non-vector-borne routes, for instance-sexual, blood transfusion, and transplacental [1]. This trait is not common in related flaviviruses. Also, ZIKV infection typically causes mild clinical symptoms; however, severe neurological complications arise post-infection [2]. Its genome translates into a single polyprotein in the following order: “C-prM-E-NS1-NS2A-NS2B-NS3-NS4A-NS4B-NS5”[3,4]. This polypeptide chain constitutes all ZIKV proteins, where the first three (Capsid, pre-Membrane, Envelope) play structural role in virus particle formation while the remaining of them (NS 1-5) are majorly involved in replication of the viral genomic RNA (vgRNA). The proteome is composed of ordered and intrinsically disordered protein regions[4–7] During translation, which takes place close to the endoplasmic reticulum (ER), the polyprotein is threaded back and forth through it (ER) with the help of multiple transmembrane domains (TMD) [3]. Consequently, topological distribution of the proteins remains exposed to either the cytosolic side or to the ER lumen. The 104 amino acid long capsid (C) protein is the first to be translated and remains attached to the ER at its C-terminal end using capsid anchor (18 amino long) while the N-terminal region faces the cytosol. Based on cryo-EM study, where they have taken 122 residues including capsid anchor there are five-helices [4,8,9]. From this study, the capsid anchor is the fifth alpha helix. The-5, also known as Capsid anchor (CA) is a single-pass transmembrane region that anchors C to the ER and separates it from the neighboring prM protein. During polyprotein processing/maturation, the viral NS2B-NS3 protease cleaves at the C-ER region, releasing the mature C protein into the cytoplasm. On the other hand, CA-prM junction on ER lumen, serves as a cleavage site identified by host signal peptidase, thus eventually separating the membrane-embedded CA from both the proteins [10,11].

ZIKV displays neurotropism, and its infection, therefore, -, and congenital Zika syndrome [12–15]. Other cell types are also highly permissive for ZIKV replication where this virus dysregulates cell cycle and cause cell death [16,17]. The underlying fundamental mechanism involved remains elusive. Despite a growing body of evidence about protein/peptide misfolding and self-assembly into pathogenic amyloids, this phenomenon in flaviviral infection is yet to be recognized [18,19]. Moreover, several diseases such as alzheimers, parkinsons, cystic fibrosis, retinitis pigmentosa, hypercholesterolemia, diabetes insipidus, and hypogonadotropic hypogonadism are linked to misfolded transmembrane domains of proteins [20]. Using a combination of bioinformatics prediction tools, fluorescent dye binding assays and high-resolution microscopy we showed that CA formed typical amyloids with short and long interconnected fibrils. Finally, our MTT and hemolysis assay further demonstrates that the CA exerts toxicity and membrane damage. All together, based on our findings we propose a new role of CA as amyloid fibrils in ZIKV pathogenesis. Considering Zika is relatively new virus among the flavivirus family, an in-depth know-how of the infection mechanism and their molecular determinants is therefore urgently needed to tackle this rapidly evolving virus. In that direction, our results therefore highlights another probable mechanism, which should be further explored to discover a new paradigm in virus-host relation and infections.

## Material and Methods

### 1. In silico analysis of capsid anchor’s aggregation propensity profile

The Zika virus (Mr766 strain; Accession no. Q32ZE1) CA peptide sequence: “GADTSIGIIGLLLTTAMA” was used to predict the aggregation propensity and amyloidogenic hotsspot using four different prediction tools, namely TANGO [21], Aggrescan [22], CamSol [23], and FoldAmyloid [24]. The sequence information was fed first and then parameters were set to run the software. Briefly, in TANGO, the parameters used were that of physiological condition (pH=7.4; temp=37°C) except for the ionic strength, which was left to default. In Camsol, Aggrescan, and FoldAmyloid complete default parameters were used. The physicochemical property and hydrophobicity index for CA was calculated using PROTPARAM (www.expasy.org/protparam) and Peptide2.0 (www.peptide2.com□) webserver tools.

### 2. Capsid anchor aggregate preparation

The CA peptide were obtained (Thermo fisher Scientific) and stored at −20°C in lyophilized form. Prior to performing any experiments, the peptides were first equilibrated to room temperature to prevent condensation and then weighed in sterile eppendorf tubes. Since CA is highly hydrophobic in nature, 1,1,1,3,3,3-hexafluoro-2-propanol (HFIP) solution was used to reconstitute the peptides at 1mg/ml concentration. The solution was then pipetted very carefully to remove any pre-formed aggregates or clumps and then allowed to sit in desiccator for overnight to evaporate off HFIP. The aggregation of ZIKV CA was initiated by dissolving peptide in 20 mM phosphate buffer (pH 7.4) and subjected to constant shaking at 1200 rpm and 37°C for upto 10 days (240 hrs). The final peptide concentration was maintained at 1mg/ml. Different time interval samples were collected for performing experiments.

### 3. Thioflavin T binding assay and kinetics

The degree of CA β-aggregation and its kinetics was determined using Thioflavin T (ThT) and Bis-ANS fluorescent dyes. ThT stock was prepared at 200 μM concentration in Milli-Q water and the binding assay was performed at a final concentration of 20 μM. The emission spectra were recorded from 470 to 700 nm at 450 nm excitation wavelength in TECAN Infinite M200 PRO multimode microplate reader using flat-bottomed black 96-well plate. Further, ThT kinetics during CA incubation was monitored by measuring ThT fluorescence with fixed amount of CA peptide sample (final concentration of 30 μM in 20 mM sodium phosphate buffer, pH 7.4) at a different time interval. Excitation and emission wavelength was set at 450 nm and 490 nm respectively. All readings were acquired in triplicates and the final value used for data-plot represents the average of three measurements. Error bars are standard deviation for three independent measurements.

### 4. Bis-ANS fluorescence assays

Bis-ANS binding assays were performed using the dye at a final concentration of 20 μM. CA monomer and aggregated samples were used at a final concentration of 30 μM (in 20 mM sodium phosphate buffer, pH 7.4). Sample was excited at 380 nm and emission was measured from 400 to 650 nm using the TECAN Infinite M200 PRO multimode microplate reader. Only Bis-ANS dye mixed with buffer was used as control. All the readings were acquired in triplicates and final plotted value represents the average of three measurements. Error bars denotes standard deviation for three independent measurements.

### 5. High-Resolution Transmission Electron Microscopy (HR-TEM)

The morphology of CA aggregates were observed with a high-resolution transmission electron microscope (FP 5022/22-Tecnai G2 20 S-TWIN, FEI). Sample preparation involved taking 7μl of the CA aggregated sample and mounted on 200 mesh carbon-coated copper grids (Ted Pella, Inc, USA). For negative staining, 3% ammonium molybdate solution was used following which the grids were air-dried overnight and images captured in HR-TEM with accelerating voltage at 200 kV.

### 6. Atomic Force Microscopy (AFM)

The aggregated CA sample were diluted 20 times and 10 μL sample were drop-casted onto freshly cleaved MICA sheet (Ted Pella, Inc, USA). After 1-hour of incubation, sample was rinsed with deionized water for three to four times and dried at room temperature overnight. The images were aquired using tapping-mode AFM (Dimension Icon from Bruker).

### 7. Cell Viability Assay

The effect of CA amyloid fibrils on cell viability was evaluated in SH-SY5Y human neuroblastoma cells. Cells was maintained in Dulbecco’s modified Eagle’s medium/Nutrient Mixture F-12 Ham (DMEM F-12) supplemented with 10% fetal bovine serum (FBS). The aggregated sample was re-suspended at calculated volume of DMEM to obtain the desired concentration. The cells were seeded in 96 well-plate at a cell density of 3000 cells/well and incubated for 24 hrs for allowing them to surface adhere. Cells were treated with different concentrations of CA aggregates from 1-250 μM and incubated at 37°C for 72 hrs. Further, 3- (4,5-dimethylthiazol-2-yl)-2,5-diphenyl tetrazolium bromide (MTT) was added to the culture medium with final concentration of 0.5 mg/mL and incubated for 3 hrs in a CO_2_ incubator. Then 100μl of DMSO solution was added to dissolve formazan crystals, incubated for 10 min and the absorbance for each well was recorded at 590 nm with a reference wavelength of 630 nm using TECAN Infinite M200 PRO multimode microplate reader. The data is reported as the percentage of the MTT reduced by the cells, and cell viability was shown relative to control cells which were not treated with the CA aggregates.

### 8. Hemolysis assay

The hemoglobin release in plasma due to RBCs lysis were monitord upon incubating with CA fibrils. The experimental procedure was reviewed and approved by the Institution’s (IIT Mandi) Ethical Committee (Approval letter number: IIT Mandi/IEC (H)/2020/17th January/P1). Before conducting experiments, the informed consent was obtained from a healthy volunteer donor. The fresh blood sample was obtained from health centre of IIT Mandi and RBCs were separated through centrifugation at 1500 rpm for 10 minutes and washed three times with isotonic phosphate buffered saline (20 mM PBS, 150 mM NaCl, pH 7.4). Further, a 10% v/v suspension of RBCs was prepared, and 200 μL of the suspension was incubated with an equal volume of CA fibril solution at varying concentrations (upto 50 μM) with shaking (300 rpm) at 37 ^o^C for 3 hrs. Finally, samples were centrifuged at 1600 rpm for 10 min and supernatant was collected in 96 well plate. Absorbance measurement was done at 540 nm in TECAN Infinite M200 PRO multimode microplate reader. All measurements were performed in triplicates. The hemolysis was calculated in relation to the hemolysis of RBCs with 1% Triton X 100, which was taken as 100%. RBCs treated with the PBS were used as control. Finally, hemolytic activity of the CA fibrils was determined in percentage using following Equation.

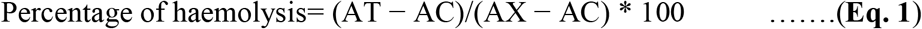

Where, AT represents the absorbance of treated supernatant, AC is the absorbance of PBS treated supernatant, and AX represents the absorbance of 1% Triton X 100 lysed cell supernatant.

## Results and Discussion

### 1. In-silico analysis revealed high aggregation propensity of ZIKV Capsid anchor

Numerous proteopathic disorders like neurodegenerative, cardiovascular, prion disease and diabetes are associated with protein misfolding and aggregation [18,19]. Typically, such disease causal proteins/peptides possess inherent aggregation prone regions (APRs) within their sequence [25]. APR’s predisposes them to oligomerize into insoluble precursor proto-fibrils that eventually give rise to amyloid fibrils [26,27]. Extensive studies on these proteins have led to recognition of common properties and structural features [28]. Thus, APR’s of any given polypeptide sequence can now be predicted with high precision tools/software’s available freely online [29,30]. In this study, we used four different open source computational tools, namely TANGO, Aggrescan,, CamSol, and FoldAmyloid to predict the aggregate forming propensity of ZIKV’s CA.

Our results revealed very high prediction score for aggregation propensity of CA, especially for residues between 7^th^-15^th^ (**Figure 2 A-D**). Typically, membrane proteins or transmembrane regions are more susceptible to self-association and aggregation because they are highly hydrophobic in nature [31]. Several predictive models developed based on membrane protein property, indicate a comparatively higher aggregation propensity in them [32]. The well-known amyloid-ß peptide (Aß) responsible for Alzheimer’s disease also has some parts of a proteolytic fragment of membrane protein-amyloid precursor protein (APP). When we analyzed the physicochemical parameters of our CA sequence using Protparam [33] and hydrophobicity index calculator (www.peptide2.com), we found the peptide to be dominated mainly with hydrophobic residues with a total 13 of them distributed across the entire peptide sequence length (Hydrophobicity index: 55.56%) (**Figure 1**). We further performed in-vitro studies on CA peptide at the physiological aggregation condition based on our preliminary results.

**Figure 2:**
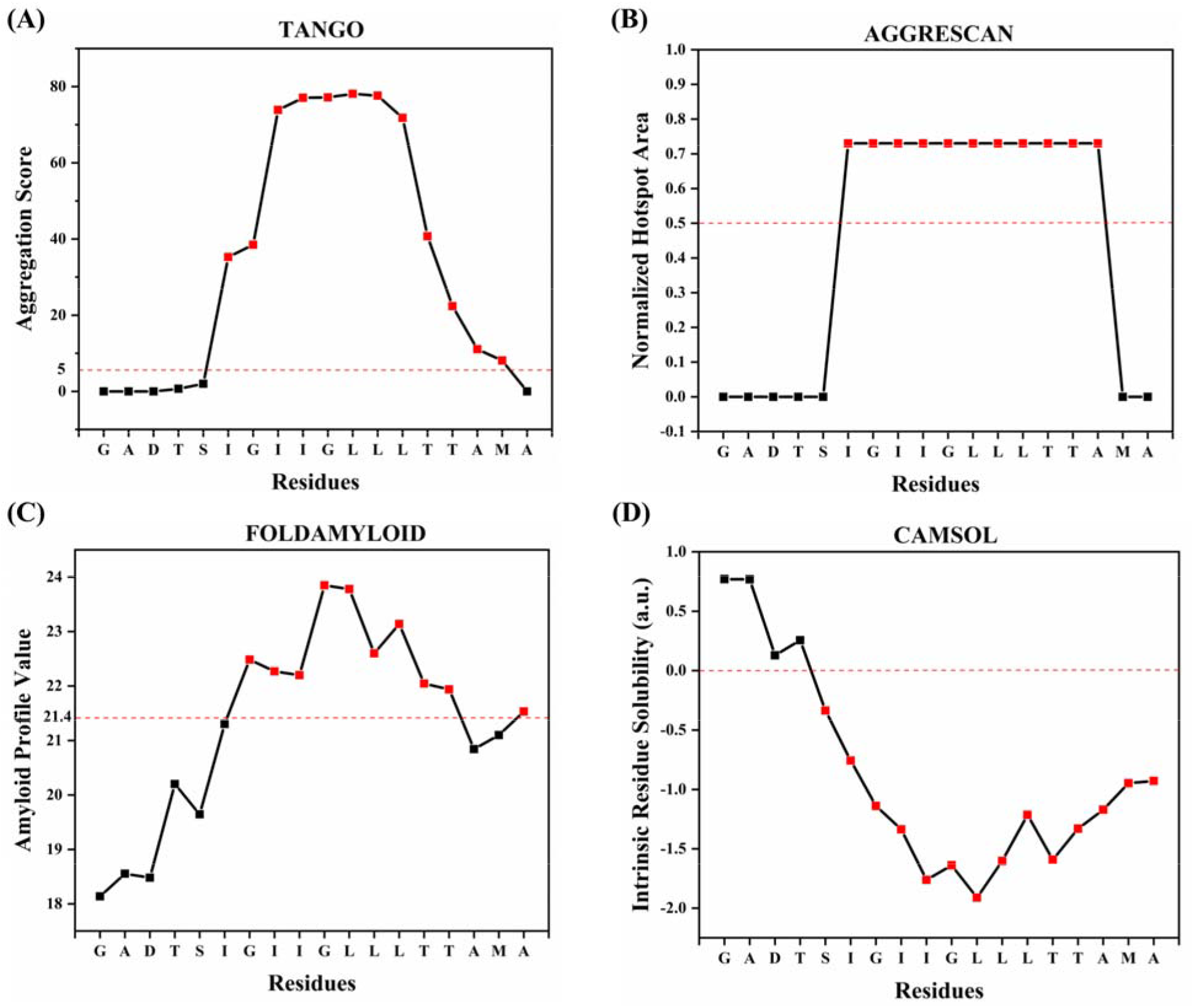
Prediction of aggregation-prone regions in Capsid anchor. APRs of Zika virus’s CA peptide sequence predicted using four different computation tools. **(A)** TANGO aggregation prediction profile indicating the percentage likelihood of a region being aggregation-prone. **(B)** Aggrescan aggregation profiles indicating the “normalised hotspot area” for APRs. **(C)** FoldAmyloid aggregation prediction indicating region as being amyloidogenic if the average value of the parameter over this region is greater than threshold (+21.4). **(D)** CamSol solubility prediction profiles with threshold values for soluble (+1) and insoluble (−1) regions shown. The X-axis in plots A, B, C represents aggregation probability with higher values corresponding to higher propensity, except for plot D where it denotes lower solubility which corresponds to higher aggregation profile.

**Figure 1:**
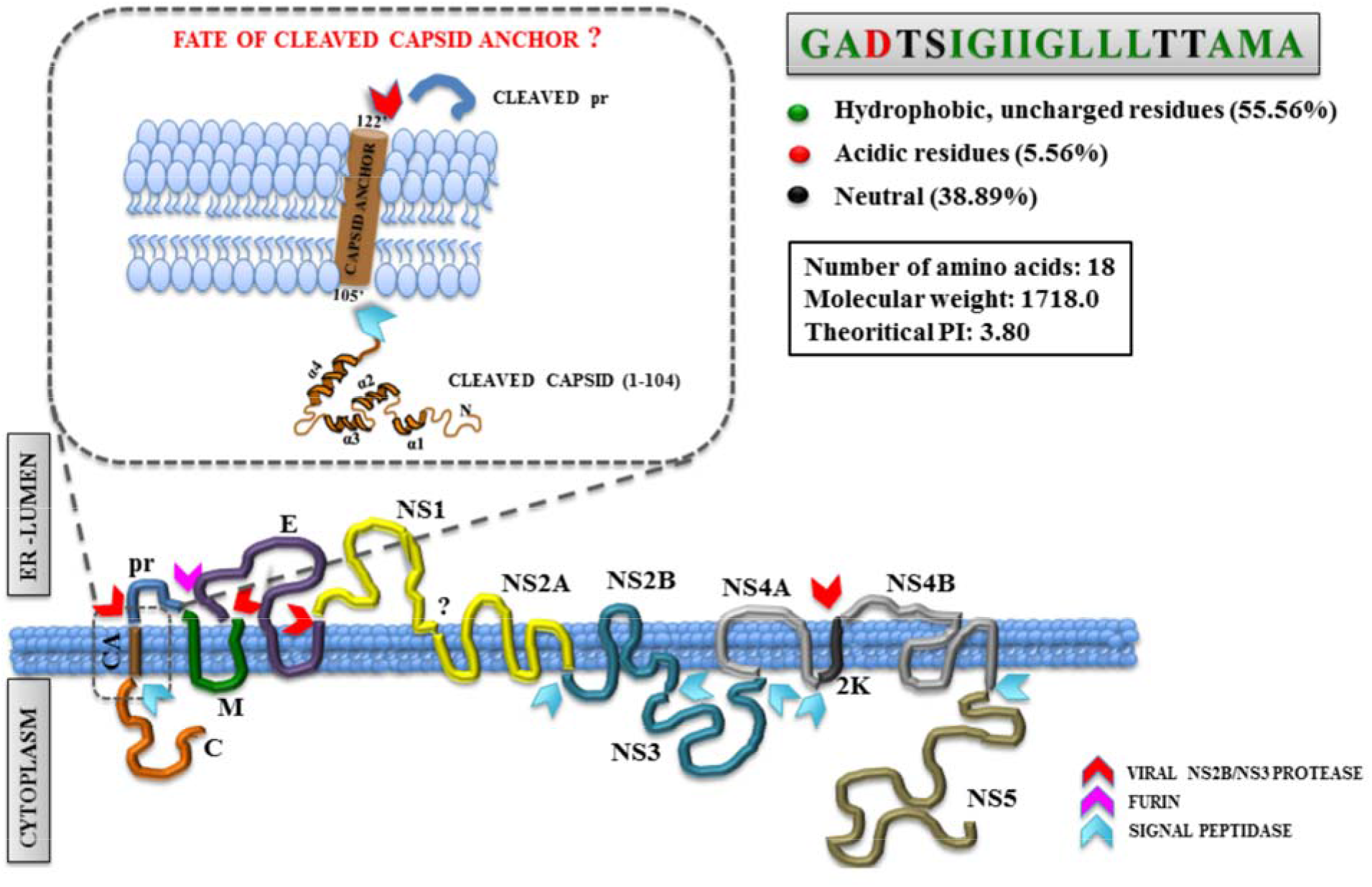
Capsid anchor topology and sequence analysis. A schematic representation of the translated polyprotein chain on ER membrane is shown. Individual proteins are represented with single-letter code, a description of which is provided down below the figure legend. The red, pink, and blue coloured arrows represent cleavage sites recognized by viral protease and host signal peptidase, respectively. Topology of membrane-embedded capsid anchor peptide is further highlighted (zoomed) post cleavage at both ends of its cytoplasmic and ER lumen site. Zika virus’s capsid anchor sequence analysis for percent hydrophobicity and physiochemical properties are indicated in respective grey and white boxes. [C=Capsid protein; CA=Capsid anchor signal peptide; pr=premembrane protein pr; M=Membrane protein; E=Envelope protein; NS1-NS5= Non-structural protein 1 to 5; 2K: signal peptide 2K]

#### 2. Capsid anchor fibrils binds with amyloid specific dyes and shows an increase in fluorescence

ThT & Bis-ANS assays are popularly used to study in vitro amyloid aggregation of proteins/peptides [33–35] ThT dye, for instance, exhibit a substantial increase in fluorescence emission upon binding to amyloid fibrils [34]. Bis-ANS, on the otherhand binds hydrophobic surfaces that result in increased fluorescence with a characteristic blue shift in emission spectra [34]. From our ThT assay, we found that ZIKV CA displays a large increase in fluorescence intensity compared to its monomeric state and control (ThT only with buffer). CA aggregates here showed ThT positive with ~15 fold increase in fluorescence emission intensity; it indicates towards the formation of amyloid fibrils upon incubation under physiological conditions.

Further, to study the amyloid formation rate by CA, we performed ThT kinetics that followed a typical sigmoidal curve with a brief lag and log phase, and a comparatively extended stationary phase (**Figure 3A**). The ZIKV CA peptide displayed a high aggregation propensity with approximately very small or no lag phase. Our supporting experiment with Bis-ANS assay further confirmed the formation of fibril by ZIKV CA peptide (**Figure 3B**). Reading acquired with sample at 0 h & 96 hrs of aggregation-inducing reaction showed an approximately 1.5-fold increase in Bis-ANS fluorescence intensity. At 240 hrs, it shows a significant increase in fluorescence with a characteristic blue shift from 549 nm to 516 nm.

**Figure 3:**
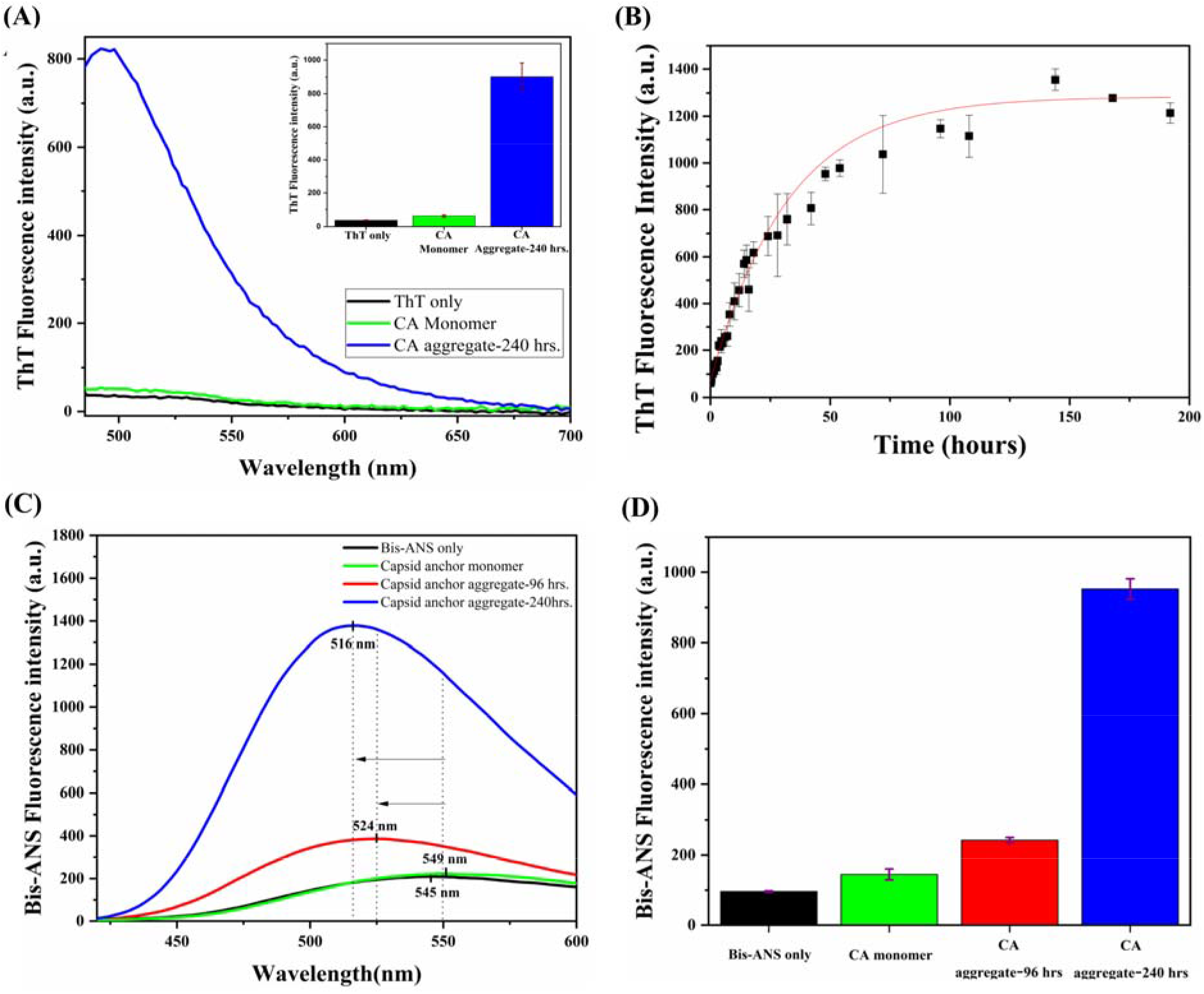
Fluorescent dye-binding assay for detection of capsid anchor aggregation. **(A)** ThT fluorescence spectra. The blue, green, and black colored lines represent spectra scan of aggregated CA, CA monomer, and ThT dye only, respectively. An increase in spectra of aggregates in comparison to its monomeric form is further displayed through bar diagram **(B)** A time-dependent ThT kinetics of CA in the presence of incubated samples. In the plot, black square represents readings of experimental data obtained at a particular time point which is fitted into a sigmoidal curve (red line). **(C-D)** Bis-ANS fluorescence spectra of aggregated CA samples at 96 hours (red), 240 hours (blue), CA monomer (green), and Bis-ANS dye only (black), respectively. The blue shift observed in the spectra scan between monomer and an aggregated sample is indicated using black dotted lines and arrows in figure (C).

#### 3. Capsid anchor aggregates display typical fibrils in TEM and AFM

In addition to monitoring dye binding properties, we then performed HR-TEM & AFM analysis to characterize and quantify the morphological features of CA aggregates. Aggregated samples of CA were acquired for TEM imaging in order to observe amyloid fibril formation. The TEM images revealed a typical amyloid structure with dense thread-like formation composed of long and short inter-connected fibrils (**Figure 4A**). The TEM images acquired in our study were analogous to previously reported amyloids produced by many cytosolic and transmembrane protein aggregates [35–37]. Further, with AFM images (**Figure 4B (I)**), we quantitatively measured the height of the CA amyloid fibrils that were obtained from the same batch previously used for TEM microscopy. We found the height of fibrils with nearly 5.5 nm (**Figure 4B(II)**).

**Figure 4:**
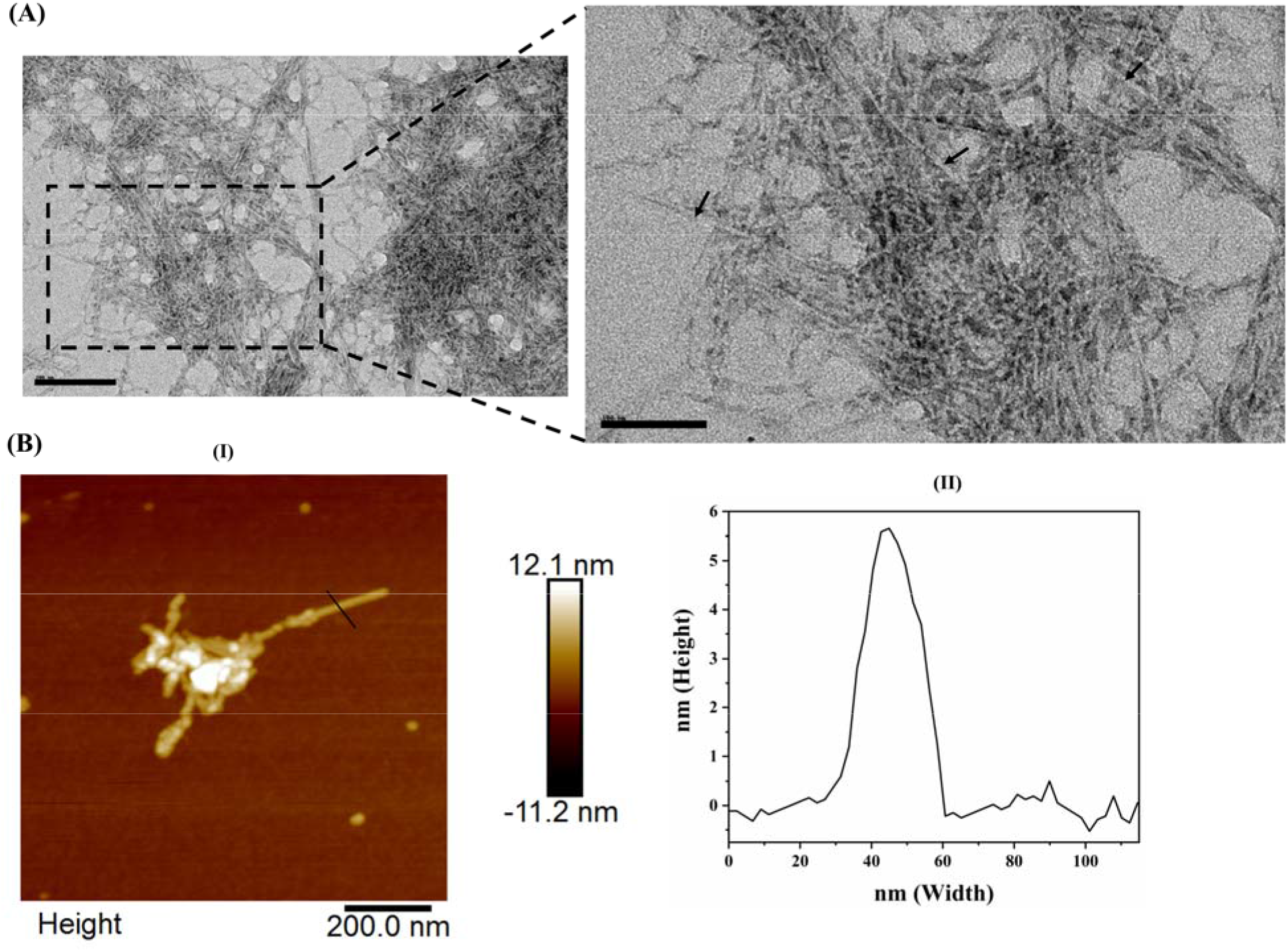
Morphology of capsid anchor amyloid fibrils. **(A)** HR-TEM images recorded at an accelerating voltage of 200 kV. Scale bar represents 200 nm and 100 nm, respectively. Black arrows in zoomed image (100nm; top right panel) are used to indicate single fibrils. **(B)** AFM image with height (Y-axis) and width (X-axis) of aggregated CA samples (I). The black line represents the fibril area selected for height measurement. The measured height of the selected fibril region is plotted (II).

#### 4. ZIKV Capsid anchor aggregates are cytotoxic amyloids

The amyloid assemblies and their cytotoxic nature has been documented in many studies [28,36,38–40]. Amyloid structures, whether as precursor oligomers or mature fibrils, possess the potential to induce membrane damage and cause cell deaths [38]. We were, therefore curious to know the effect of ZIKV CA amyloid fibrils on the viability of human cells. We used human neuroblastoma SH-SY5Y cells for our study, as this particular cell line has been used previously for studying ZIKV infection as well as amyloid-β fibril toxicity [16,37,39]. The cells were incubated with different concentrations of CA amyloid aggregates ranging from 1-250 μM concentrations and incubated for a period of 72 hrs. We observed that with increasing concentration of CA aggregates, the percent cell viability reduced, suggesting their cytotoxic nature (**Figure 5A**). Cell viability remained ~35% for the highest aggregate concentration used in comparison to control (**Figure 5A**). Having established the cytotoxic property of CA amyloid fibrils, next, we performed hemolysis assay with the same aggregate sample to illustrate if the cell death observed were membrane damage induced. Thus, RBC was isolated following standard protocols and treated with CA amyloid fibrils in a concentration-dependent manner (0.3-50 μM). Upon measurement of hemoglobin specific absorbance, we found increasing absorbance yield with higher aggregate concentration (**Figure 5B**). Up to ~4% hemolysis could be seen in 3 hrs incubation period at the highest amyloid concentration. Previously, the membrane-lytic activity of an influenza A virus protein “PB-F2” was linked to cell death induced by amyloid oligomers of the protein [40]. Our results thus clearly illustrate membrane damaging and cytotoxic nature of ZIKV CA derived aggregates/amyloid-fibrils.

**Figure 5:**
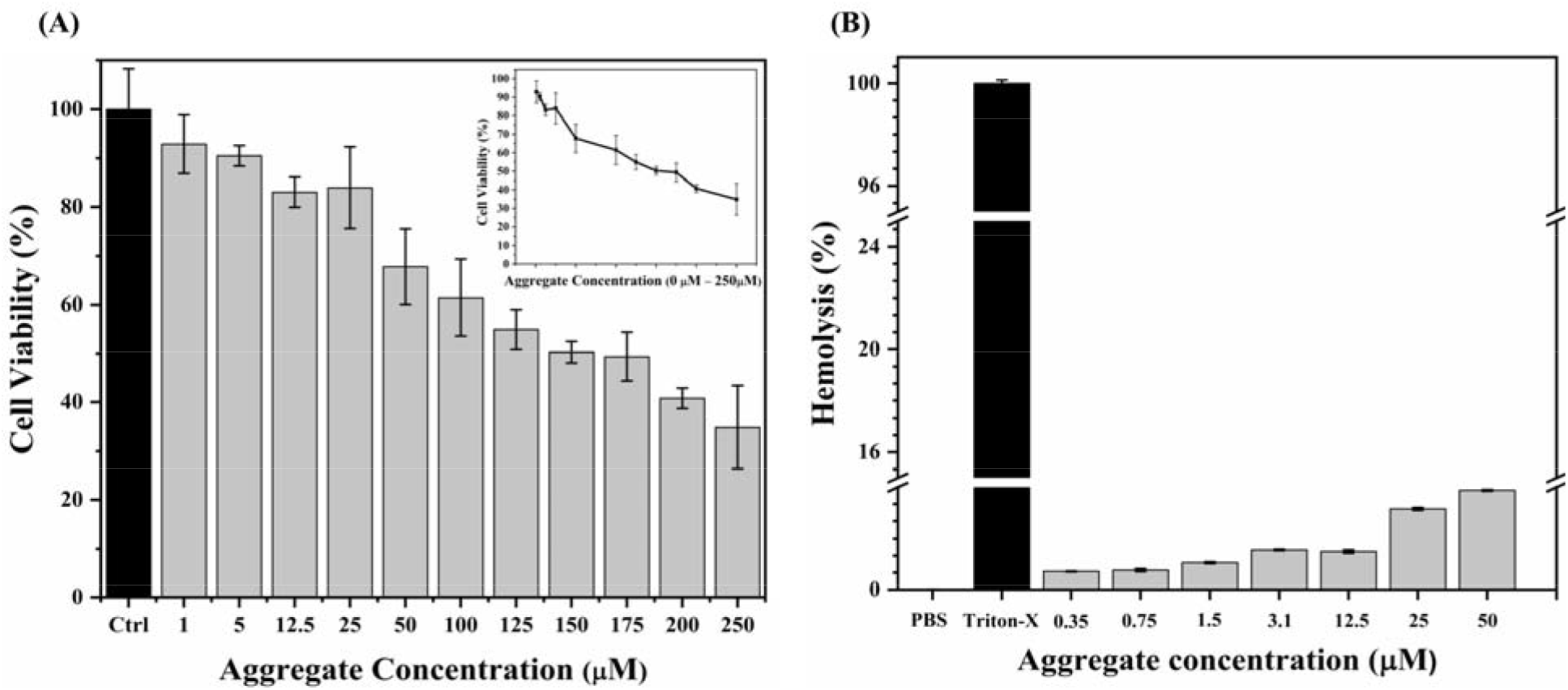
MTT assay and RBCs hemolysis assay. **(A)** The cell viability assay on SH-SY5Y cells incubated with different concentration of CA aggregates (0-250 μM) for 72 hrs. The percent cell viability (equivalent to percent reduction in MTT) showing cytotoxic nature of CA amyloid fibrils relative to PBS treated control cells is plotted. **(B)** The hemolysis assay on human blood extracted RBC’s at different concentration of CA aggregates. The bar graph represents increase in absorance of aggregate treated RBC’s against PBS treated and Triton-X treated RBC’s used as positive and negative controls.

### Discussion

Previously, studies performed to understand CA revealed their numerous roles in the viral life cycle[9,10,41-43] In addition to its canonical function of anchoring C to ER and acting as a signal sequence for prM, CA also regulates the maturation of viral proteins and their assembly into new infective particles[10,41,42]. A recent study on ZIKV’s CA explicitly highlighted some unique properties that further enhanced our understanding of this flaviviral transmembrane domain[41]. They reported that ZIKV strictly requires its own CA sequence for virus-like particle (VLP) formation, whereas DENV and TBEV can form VLP in the presence of other virus-derived heterologous CA sequences[10,41]. Also, mutation or complete replacement of ZIKV CA sequence adversely affects the stability of downstream E protein resulting in their degradation and failed viral assembly[10,11,41]. A separate study highlighted the importance of CA in trimerization of capsid dimers, which is essential for interaction of downstream proteins prM and E with their TMDs[9]. However, the role of CA, in isolation, in mediating cytopathic effect in ZIKV pathogenesis remains unexplored.

In this study we highlight yet another mechanism through which it possibly aids ZIKV induced cellular pathogenesis i.e. through cytotoxic amyloid fibril formation. We found the synthetic capsid anchor peptide readily forms typical amyloid-like fibrils as confirmed by fluorescence dye-based assays and high-resolution microscopy. These fibrils caused cell death and RBC lysis in a concentration-dependent manner. However, the exact mechanism involved needs to be studied in future. Our findings can be correlated to the neuropathological defects in ZIKV-infected individuals and paves way for exploring aggregation propensities of various other TMD’s spanning the ZIKV polyprotein. Also, the link with alzheimer’s amyloid beta would be an interesting question. In conclusion, this study provides invaluable information about the amyloidgenic nature of ZIKV’s CA peptide.

## Authors’ contribution

RG: Conception, design, review, and writing of the manuscript. KUS, KG, and AK performed experiments, analyzed data and wrote the manuscript.

## Acknowledgment

Authors are grateful to the Indian Institute of Technology Mandi (BioX, C4DFED, and AMRC center) for all the facilities. The work was supported from Department of Biotechnology (DBT), Government of India (BT/11/IYBA/2018/06) to RG. KUS is thankful to Indian Council of Medical Research (ICMR), India, for fellowship. KG was supported by the Science and Engineering Research Board (SERB), India (CRG/2019/005603). AK was supported by DBT, India (BT/11/IYBA/2018/06).

